# A candidate *HNRNPU*–*CHD2* RNA-processing axis in neurodevelopmental disease

**DOI:** 10.64898/2026.11.01.763498

**Authors:** Huan Tang, Zhen Xiao, Ruilan Wang, Isaac A. Babarinde, Songqiang Huang, Andrew P. Hutchins, Gang Ma

## Abstract

Heterozygous loss-of-function variants in *HNRNPU*, which encodes a nuclear RNA-binding scaffold protein, cause a neurodevelopmental disorder with epilepsy, intellectual disability and brain malformations, but the programs that link *HNRNPU* dosage to neuronal dysfunction remain poorly defined. We depleted *HNRNPU* in Human embryonic kidney 293T (HEK293T) cells, profiled gene expression and alternative splicing, intersected the two datasets, and asked which of the resulting genes already carried a neurodevelopmental annotation. Knockdown produced 5,643 differentially expressed genes and 21,728 high-confidence splicing events; 1,222 genes changed at both levels, and two pre-specified neurodevelopmental panel genes were among them: *CHD2* and *DYRK1A*. In K562 eCLIP experiments, 107 reproducible *HNRNPU* peaks were detected across the *CHD2* gene body, and three of the four regulated exons introduce a reading-frame shift whose premature termination codon lies far upstream of the final exon junction, predicting nonsense-mediated decay. Public K562 knockdown data reproduced the rise in *CHD2* mRNA and the same skipped-exon change, and *CHD2* depletion in the neural line SH-SY5Y altered cell-cycle, DNA-repair and neural gene programs. HNRNPU therefore constrains a frameshift-prone splicing switch in *CHD2*, a candidate route from *HNRNPU* haploinsufficiency to loss of functional *CHD2* if these transcripts are degraded.

**Highlights:**

- *HNRNPU* depletion changes *CHD2* expression and exon inclusion in human cells.
- eCLIP places *HNRNPU* on *CHD2* RNA, consistent with direct control of its splicing.
- Three *CHD2* exons are frameshift-prone, predicting loss of full-length *CHD2*.

## Introduction

*HNRNPU* encodes heterogeneous nuclear ribonucleoprotein U (also known as *SAF-A*), an abundant nuclear protein that contains an N-terminal SAP domain, which binds scaffold-attachment regions and chromatin ^[1, 2]^, and a C-terminal RGG/RNA-binding domain. *HNRNPU* oligomerizes on chromatin-associated RNA and contributes to the establishment of higher-order interphase chromosome structure ^[3–5]^. In our previously published work, we showed that *HNRNPU* anchored the promoters of primed-specific genes to the nuclear matrix, promoted their expression and RNA stability, and that its depletion destabilized the primed state in human pluripotent stem cells ^[6]^. In addition to its architectural role, *HNRNPU* acts on RNA metabolism, it associates with the spliceosome and has been reported to exert global control over alternative splicing through regulation of U2 snRNP maturation ^[7]^.

Heterozygous loss-of-function variants in *HNRNPU* cause a recognizable neurodevelopmental disorder with a consistent core phenotype, comprising early-onset epilepsy, severe intellectual disability, absent or delayed speech, hypotonia, hypoplasia of the corpus callosum and dysmorphic features, but with variable severity between individuals ^[8, 9]^. Mouse models show that *HNRNPU* is required for cortical development and that reducing its dosage produces seizures and behavioral deficits ^[10, 11]^, so the human condition is best explained by reduced dosage rather than by absence of the protein. The mechanism by which lower *HNRNPU* dosage produces this phenotype is unresolved, a difficulty that is typical of neurodevelopmental genetics more broadly, where a molecular diagnosis often does not yet explain how the affected gene produces disease ^[12]^. Because *HNRNPU* is a broadly expressed nuclear factor with roles in both chromatin architecture and RNA processing, potential mechanisms range from global transcriptional dysregulation to selective effects on a small number of dosage-sensitive targets ^[10, 11]^.

*CHD2* encodes chromodomain-helicase-DNA-binding protein 2, an ATP-dependent chromatin remodeler of the SNF2 family that is highly expressed in the developing brain and is required for cortical neurogenesis ^[13, 14]^. De novo loss-of-function variants in *CHD2* cause developmental and epileptic encephalopathy, including phenotypes overlapping Dravet syndrome and Lennox-Gastaut syndrome, and *CHD2* is also among the genes recurrently mutated in autism spectrum disorder ^[15–18]^. The phenotypic overlap between *HNRNPU*- and *CHD2*-related disease, comprising epilepsy with developmental delay and a substantial burden of behavioral and language impairment, led us to examine this pair of genes: both encode nuclear regulators with roles in chromatin and RNA, and both are dosage-sensitive in the developing brain. We therefore tested directly whether *HNRNPU* regulates the expression and RNA processing of *CHD2*. Whether *HNRNPU* regulates *CHD2* has, to our knowledge, not been examined.

Whether the resulting transcripts still yield functional CHD2 protein is a separate question: a factor acting on RNA processing can alter the isoform composition of a target without changing its total abundance. We therefore analyzed differential expression and differential splicing together, and asked which transcripts were affected at both levels. The same logic has been used to link RNA-binding proteins to aberrant splicing in other settings ^[19]^, and RNA-binding proteins themselves are increasingly recognized as disease genes across organ systems ^[20]^. This combined view is particularly relevant to neurodevelopmental disease, in which the genes most frequently mutated are enriched for regulators of RNA processing and chromatin ^[21, 22]^.

Alternative splicing is especially consequential in the nervous system, where it expands the repertoire of protein isoforms that shape neuronal differentiation, synapse formation and excitability. Transcript diversity also arises from the incorporation of transposable-element-derived sequences into coding and non-coding transcripts, which can modulate the transcriptome ^[23]^. The regulators of these decisions are themselves enriched among the genes mutated in neurodevelopmental disorders ^[21, 22]^, and reducing the abundance of an RNA-binding protein typically shifts the inclusion of many exons rather than the abundance of a few transcripts ^[7, 19]^. Such shifts can be pathogenic without any change in total mRNA: skipping of a cassette exon may delete a protein domain, disrupt an interaction surface, or introduce a premature termination codon that targets the transcript for nonsense-mediated decay. For a dosage-sensitive regulator, this creates a second route to disease in which the damaging event is the loss of a specific isoform rather than the loss of the gene. Isoform-level consequences of this kind are invisible to expression-only analyses, which is why we quantified transcript abundance and exon inclusion together and then translated the affected isoforms.

We therefore depleted *HNRNPU* in HEK293T cells with two independent shRNAs, integrated differential expression with differential splicing, tested the regulated exons against public *HNRNPU* eCLIP occupancy, and translated the affected isoforms to determine whether the transcripts could encode a functional protein. We then depleted *CHD2* in the neural line SH-SY5Y to ask which programs depend on this node. Our data support a model in which *HNRNPU* occupies *CHD2* pre-mRNA and constrains a cassette-exon switch whose products are truncated before the catalytic domains, linking *HNRNPU* dosage to a neurodevelopmental gene program.

## Materials and Methods

### Study design

The study had three stages (**Figure 1**). In the discovery stage, *HNRNPU* was depleted in HEK293T cells with two independent shRNAs and the cells were profiled by RNA sequencing; differential expression and differential alternative splicing were then analyzed jointly, so that genes altered at both levels could be identified. In the prioritization and mechanism stage, this intersection was interrogated with a panel of 32 neurodevelopmental genes, and *CHD2* was tested against public HNRNPU eCLIP data and analyzed for reading-frame and nonsense-mediated decay (NMD) consequences. In the functional stage, *CHD2* was depleted in the neural cell line SH-SY5Y to ask whether its loss affects neural and cell-cycle programs; public HNRNPU knockdown datasets were analyzed in parallel to place the findings in context.

**Figure 1.**
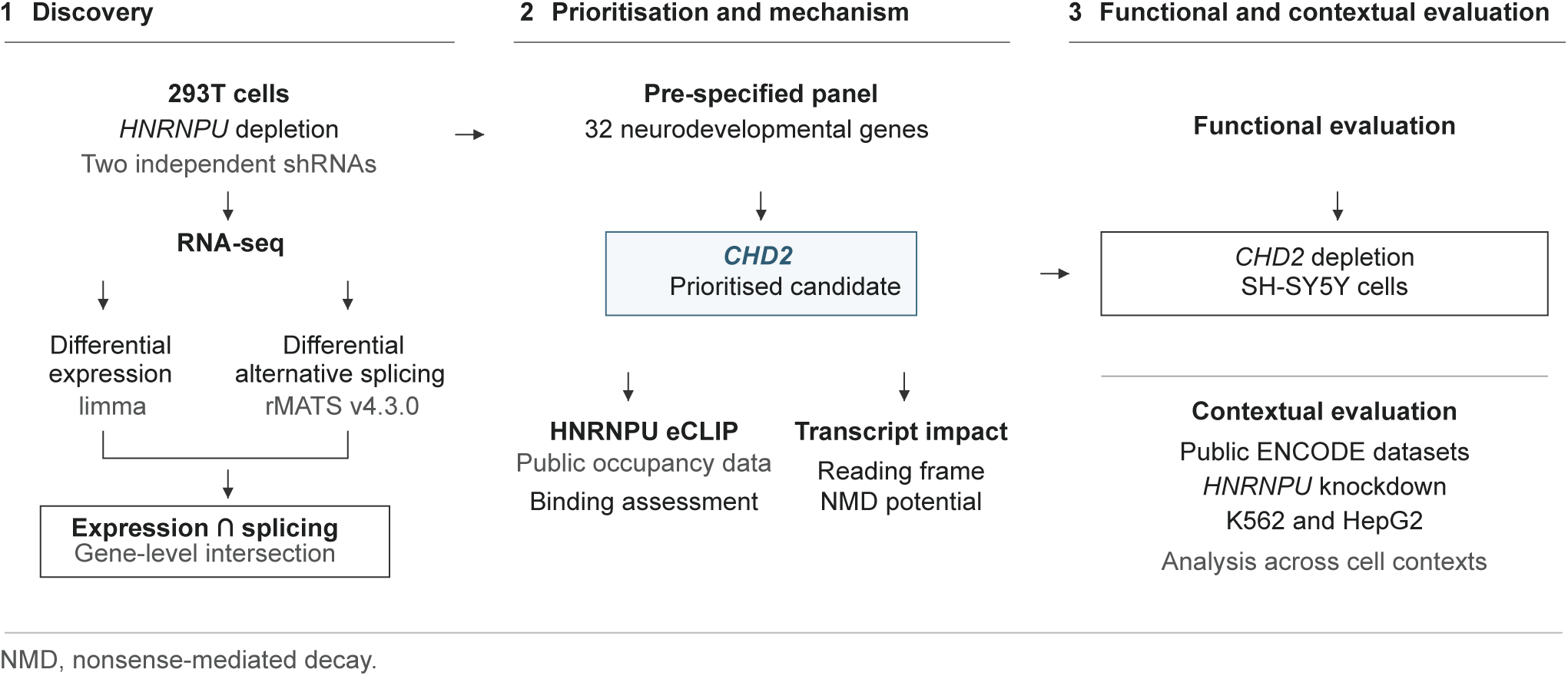
Experimental and analytical design. The study comprises three stages. Stage 1 (discovery): *HNRNPU* was depleted in HEK293T cells with two independent shRNAs, and RNA sequencing data were analyzed for differential expression (limma) and differential alternative splicing (rMATS v4.3.0); the two outputs were intersected. Stage 2 (prioritization and mechanism): the expression × splicing intersection was interrogated with a pre-specified 32-gene neurodevelopmental panel, and the resulting candidate was tested against public *HNRNPU* eCLIP occupancy data and evaluated for reading-frame and nonsense-mediated decay consequences. Stage 3 (functional and contextual evaluation): *CHD2* was depleted in SH-SY5Y cells, and public ENCODE *HNRNPU* knockdown datasets in K562 and HepG2 were analyzed for context.

### Cell culture and knockdown

HEK293T cells were used for the *HNRNPU* knockdown discovery experiment, and the human neuroblastoma line SH-SY5Y was used for the *CHD2* knockdown experiment. Both lines were maintained under standard conditions in Dulbecco’s modified Eagle’s medium supplemented with fetal bovine serum at 37 degrees Celsius and 5% carbon dioxide. Knockdown was achieved by lentiviral delivery of shRNAs, with a non-targeting shRNA against luciferase (sh*LUC*) as the control in all experiments. Two independent *HNRNPU* shRNAs (shHNRNPU-1 and shHNRNPU-2), which have been shown to reduce *HNRNPU* at both the RNA and protein level ^[6]^, and two independent effective *CHD2* shRNAs were used. Knockdown efficiency was assessed from transcript abundance in the sequencing data, and growth was monitored as the cumulative population doubling number over four serial passages.

### RNA sequencing and read processing

Total RNA was extracted and sequenced by a commercial provider. Reads were aligned to the human reference genome (GRCh38/hg38) with STAR ^[24]^ using GENCODE v49 annotation, and gene-level counts were generated with featureCounts ^[25]^. Junction counts for the splicing analysis were obtained from the same alignments in junction-count mode. All analyses were performed against the GENCODE v49 transcript set, so that exon identifiers used in the splicing results map directly onto the canonical *CHD2*-201 model described below. The HEK293T experiment comprised two biological replicates each of sh*LUC*, sh*HNRNPU*-1 and sh*HNRNPU*-2, and the SH-SY5Y experiment comprised two biological replicates each of sh*LUC*, sh*CHD2*-1 and sh*CHD2*-2.

### Differential expression analysis

For the HEK293T *HNRNPU* experiment, the two *HNRNPU* shRNAs were analyzed jointly as a merged knockdown, which reduces the influence of shRNA-specific off-target effects; differential expression was assessed with limma 3.66.0 under R 4.5.2 ^[26]^. Merging was implemented by treating the four *HNRNPU*-depleted libraries as one group and the two control libraries as the other, which yields a single estimate of the effect of *HNRNPU* loss that is not dominated by either shRNA. Genes were considered differentially expressed at an adjusted *P* value < 0.05 and an absolute log2 fold change greater than 1; results are reported as unique gene symbols (5,643 genes, obtained by collapsing 5,646 Ensembl gene identifiers to unique gene symbols). For the SH-SY5Y *CHD2* experiment, differential expression was assessed with DESeq2 1.46.0 ^[27]^ for each shRNA individually, for the merged sh*CHD2*-1 plus sh*CHD2*-2 comparison. Genes were considered significant at an adjusted *P* < 0.05 and an absolute log2 fold change greater than 1. Core consistent genes were defined as genes that were significant with the same direction of change under both effective shRNAs.

### Differential splicing analysis

Alternative splicing was quantified with rMATS v4.3.0 ^[28]^ in junction-count mode across the five standard event classes (skipped exon, mutually exclusive exons, retained intron, alternative 5’ splice site and alternative 3’ splice site). Events were considered high confidence at an FDR below 0.05 and an absolute dPSI of at least 0.1; the strict event counts reported here (13,627 skipped-exon events out of 153,388 tested) use these thresholds. Inclusion levels (PSI) for individual events are reported as the mean of the replicate values provided by rMATS.

### Integration of expression and splicing

Genes were assigned to four groups according to the direction of their expression change (up or down) and the direction of their mean inclusion change (inclusion up or inclusion down), using an absolute log2 fold change greater than 1 and an absolute mean dPSI of at least 0.1 as thresholds. Two intersections were computed. Without restricting the event class, 1,535 differentially expressed genes carried at least one high-confidence event of any class; restricting the splicing side to skipped-exon events, the class that dominates the data, gives 1,222 genes. The skipped-exon-specific set is the one used throughout, because the candidate gene prioritized below is an exon-skipping target.

### Functional, disease and panel enrichment

Gene ontology and KEGG enrichment were performed with clusterProfiler 4.18.4 and org.Hs.eg.db 3.22.0 ^[29]^, and gene set enrichment analysis used the Hallmark, C2 and C5 collections with fgsea 1.36.2 ^[30]^. Enrichment was computed separately for each expression × splicing group, so that a pathway appearing in one group could be compared with its significance in the others; the background for each test was the full set of detected genes rather than the group itself, which prevents the appearance of spurious enrichment in small groups. Disease enrichment used DisGeNET ^[31]^ and Enrichr ^[32, 33]^ against the same background, with an adjusted *P* < 0.05 threshold, and the member genes of each enriched term were inspected for the presence of *HNRNPU*, *CHD2* and *DYRK1A*. A pre-specified panel of 32 genes implicated in neurodevelopmental disorders, epilepsy or neuronal differentiation was used to interrogate the expression × splicing intersection.

### Off-target sequence analysis

To determine whether the genes identified in the *CHD2* knockdown experiment could arise from seed-mediated off-target repression rather than on-target *CHD2* depletion, we extracted the 6-nucleotide and 7-nucleotide seed sequences of each shRNA and matched them against the 3’ untranslated regions of the detected transcripts, following the standard seed-matching approach for RNA interference specificity. For each shRNA we then tested whether the core consistent gene set, or the gene set unique to that shRNA, was enriched for seed-matched transcripts relative to the full detected transcript universe, using Fisher’s exact test with Benjamini-Hochberg correction across the twelve tests.

### Analysis of public eCLIP and knockdown data

Public data were obtained from the ENCODE portal ^[34, 35]^; the datasets, replicates and accessions are summarized in **Table S7**. *HNRNPU* eCLIP peaks generated in K562 cells (experiment ENCSR520BZQ; files ENCFF408HQB and ENCFF638WUB, GRCh38) were intersected with the *CHD2*-201 transcription unit (chr15: 92,900,324-93,027,996) and with the exons and flanking introns of the regulated cassette exons, following the analysis framework established for eCLIP data ^[36]^. Peaks were counted when they overlapped the transcription unit in each duplicate. The two peak files are the duplicate calls of a single eCLIP experiment; a peak was treated as reproducible when it overlapped a peak in the other duplicate by at least one base pair, and the reproducible count reported is the number of calls in duplicate 1 that overlapped a call in duplicate 2. Regulated exons were classified from the rMATS coordinates of the HEK293T experiment.

For expression-level comparison, gene quantification files were obtained for the K562 *HNRNPU* shRNA experiment (ENCSR047IUS) and its matched non-targeting control (ENCSR438MDN), and TPM values for *HNRNPU* and *CHD2* were extracted for all four knockdown and four control quantification sets (two biological replicates in two quantification pipelines). Differential expression files for the K562 (ENCSR047IUS) and HepG2 (ENCSR308IKH) experiments were used for the corresponding splicing comparisons. Corresponding exons between hg19 and GRCh38 were matched using the local coordinate offset in the *CHD2* 3’ region (543,230 bp), which was verified against two independent exon pairs.

### Exon model, reading frame and NMD prediction

The canonical *CHD2*-201 exon model was derived from GENCODE v49 annotation and comprises 39 canonical exons spanning chr15: 92,900,324-93,027,996 (GRCh38). Regulated cassette exons were assigned to this model by coordinate intersection. For each regulated exon, the reading-frame consequence of skipping was first predicted from the exon length modulo three and then verified by translating the reconstructed isoform, as described below. Frame shift events were then evaluated against the 50-55 nucleotide rule for NMD ^[37]^: because mature mammalian transcripts are spliced, a premature termination codon located more than 50-55 nucleotides upstream of the last exon-exon junction is expected to trigger NMD. The number of downstream canonical exons and the distance to the annotated termination codon were computed for each event.

### Co-expression analysis in the developing brain

Co-expression with *HNRNPU* was computed from the BrainSpan exon-level RPKM matrix (524 samples, 8 post-conceptional weeks to 40 years) ^[38]^. Values were log2(RPKM + 1) transformed, genes with a mean log2 value above 1 were retained (16,256 genes), and Pearson correlation with *HNRNPU* was computed across all samples; a gene’s percentile is its rank within that set (**Table S8**). The comparison is descriptive: co-expression in bulk tissue can reflect shared cellular composition or expression breadth rather than regulation, and *ACTB* is included only as a widely expressed reference point.

### ClinVar intersection

Short variants reported for *CHD2* were retrieved from ClinVar ^[39]^ through the NCBI E-utilities, and the 1,968 records carrying a GRCh38 position were intersected with the four regulated exons, their core splice sites and 2, 10 and 50 bp of flanking sequence.

### Isoform translation and nonsense-mediated decay analysis

To determine the protein-level consequence of the regulated exons we translated the affected isoforms rather than relying on exon length alone. The MANE Select transcript of *CHD2* (*CHD2*-201, ENST00000394196) and the genomic sequence of the four regulated exons were obtained from Ensembl release 113 through the REST API. The coding sequence was reconstructed with each canonical exon removed or with the alternative exon inserted at its annotated position, and the resulting open reading frames were translated. A stop codon was considered premature when it was not the terminal codon of the reconstructed coding sequence, and the distance from that codon to the last exon-exon junction was compared with the 50-55 nucleotide rule for nonsense-mediated decay ^[37]^.

### Statistical analysis

Differential expression used an adjusted *P* < 0.05 with an absolute log2 fold change greater than 1, and differential splicing used an FDR below 0.05 with an absolute dPSI of at least 0.1. Enrichment analyses used an adjusted *P* < 0.05. Comparisons of gene-set overlap between the *HNRNPU* and *CHD2* knockdown experiments used Fisher’s exact test, as did the off-target seed tests. Unless stated otherwise, adjusted *P* values are corrected using the Benjamini-Hochberg method. Biological replicates are as described above and are indicated in the figure legends.

Software versions were as follows: R 4.5.2 with limma 3.66.0, clusterProfiler 4.18.4, fgsea 1.36.2, org.Hs.eg.db 3.22.0 and enrichplot 1.30.5 for the expression, pathway and gene set analyses; R 4.5.2 with DESeq2 1.46.0 for the *CHD2* knockdown differential expression analysis; rMATS v4.3.0 for splicing quantification; and STAR with featureCounts 2.0.6 (Subread) for alignment and counting. Python 3.11.7 with pandas 2.1.4, numpy 1.26.4, scipy 1.11.4 and matplotlib 3.8.0 was used for data integration, eCLIP peak mapping and figure generation. Analysis scripts and derived tables are provided with the manuscript so that the reported values can be regenerated from the deposited counts.

## Results

### *HNRNPU* depletion reprograms the HEK293T transcriptome

We first asked whether loss of *HNRNPU* affects cell growth and gene expression in human cells. *HNRNPU* was depleted in HEK293T cells with two independent shRNAs described previously ^[6]^, using a non-targeting shRNA against luciferase (sh*LUC*) as the control. Cultures depleted of *HNRNPU* proliferated far more slowly than sh*LUC* controls: the depleted wells were less dense at the end of the experiment (**Figure 2a**), and after four serial passages the cumulative population doubling number was 8.2 for sh*HNRNPU*-1 and 5.4 for sh*HNRNPU*-2, compared with 33.3 for sh*LUC* (**Figure 2b**).

**Figure 2.**
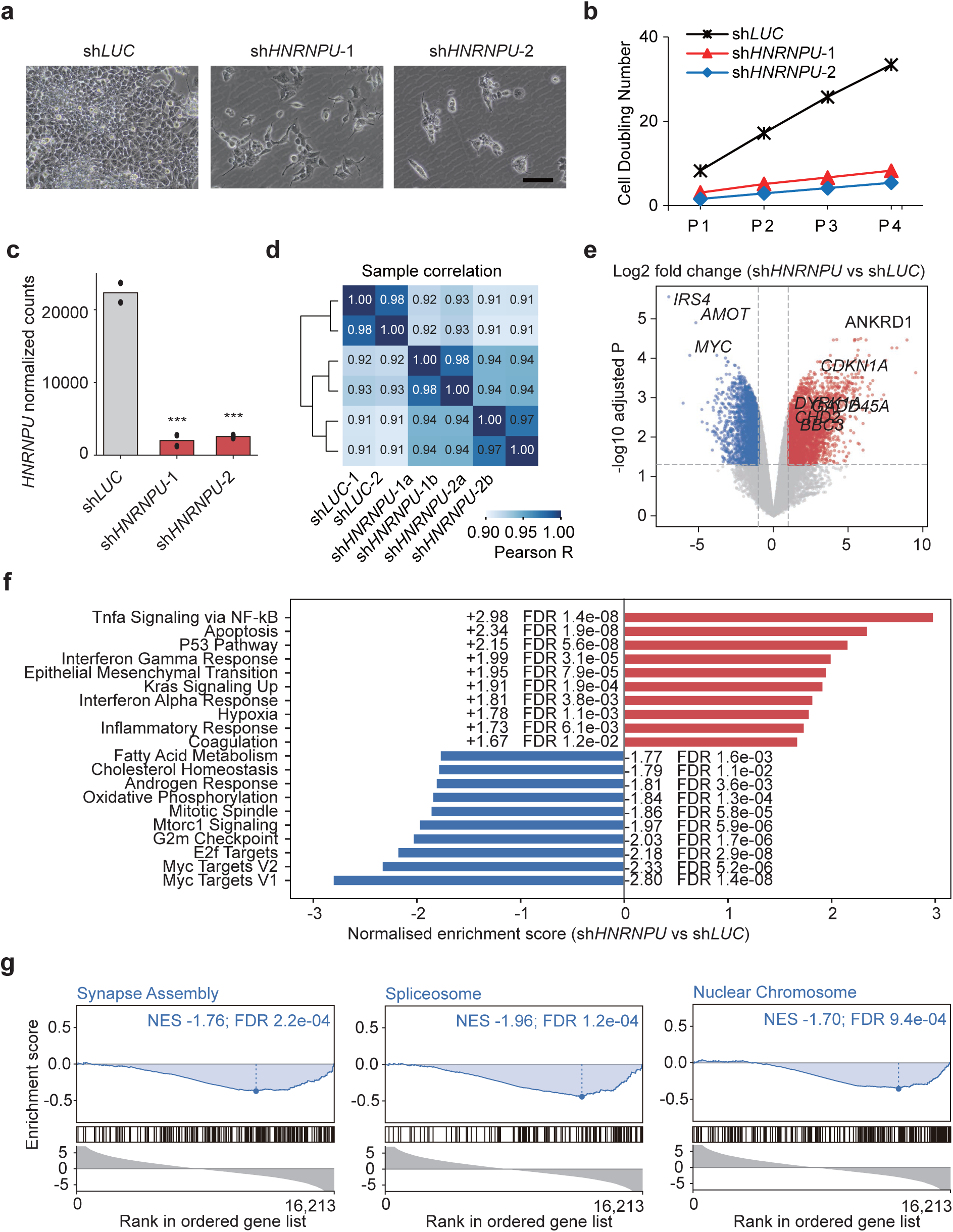
*HNRNPU* depletion impairs proliferation and reprograms the HEK293T transcriptome. a. Representative images of shLUC, shHNRNPU-1 and shHNRNPU-2 cells. Scale bar, 20 µm. b. Cumulative population doubling number over four serial passages (P1-P4) for shLUC, shHNRNPU-1 and shHNRNPU-2. c. *HNRNPU* normalized counts in the six libraries (two shLUC, two shHNRNPU-1 and two shHNRNPU-2); asterisks mark the comparison of each shRNA with shLUC. d. Sample-correlation heatmap of the six libraries (Pearson r of log2 normalized counts across all 27,114 detected genes; the color scale spans r = 0.9-1.0). Replicates of the same condition correlate more closely with each other than with samples of the other condition, which is the reason the two shRNAs were analyzed jointly rather than separately. e. Volcano plot of the merged knockdown (3,171 genes up and 2,472 down); selected genes are labeled. f. Gene sets representing cellular programs previously associated with *HNRNPU*, quantified by gene set enrichment analysis in the merged knockdown data; normalized enrichment score and adjusted P are given to the right of each set. g. Running enrichment curves for three representative gene sets (synapse assembly, spliceosome and nuclear chromosome); each panel shows the running enrichment score, the positions of the gene-set members in the ranked list (hit ticks) and the ranking metric below, with normalized enrichment score and adjusted P above.

Because a proliferation defect can arise from many different programs, we profiled the knockdown transcriptome rather than a single pathway, quantifying transcript abundance and alternative splicing together with RNA sequencing of two biological replicates each of sh*LUC*, sh*HNRNPU*-1 and sh*HNRNPU*-2, giving six libraries in total. Both shRNAs were effective in these libraries, reducing *HNRNPU* mRNA to 8.9% and 11.4% of the sh*LUC* level (**Figure 2c**). Because single-shRNA experiments can be dominated by off-target effects, we analyzed the two shRNAs jointly as a merged knockdown. Sample correlation supported this choice (**Figure 2d**): the three replicate pairs correlated at Pearson r = 0.97 to 0.98, whereas correlations between control and knockdown samples were lower (0.91 to 0.93). In the merged comparison 5,643 genes were differentially expressed at an adjusted *P* < 0.05 with an absolute log2 fold change greater than 1 (3,171 up and 2,472 down; **Figure 2e and Table S1**).

To connect these changes to previously reported phenotypes of *HNRNPU* loss of function, we converted known *HNRNPU*-associated cellular programs into gene sets and quantified them in the merged knockdown data. Proliferation and cell-cycle programs were coordinately suppressed (MYC targets V1, normalized enrichment score (NES) −2.80; MYC targets V2, −2.33; E2F targets, −2.18; G2/M checkpoint, −2.03), whereas p53, apoptosis and inflammatory programs were induced (TNFA signaling via NF-κB, +2.98; apoptosis, +2.34; p53 pathway, +2.15; all adjusted *P* < 1 × 10^-5^; **Figure 2f**). Because both neurodevelopmental phenotype and RNA processing were central to this study, we highlighted three gene sets from these themes in the running enrichment curves: synapse assembly (normalized enrichment score −1.76), the spliceosome (−1.96) and the nuclear chromosome (−1.70) were all depleted (**Figure 2g**). These gene sets were chosen for their relevance to the reported phenotype rather than selected from an unbiased ranking, and the suppression of proliferation and cell-cycle programs in particular is expected in cultures that have stopped growing, so the enrichment results indicate which programs move with *HNRNPU* dosage rather than which ones are direct targets. Together, these data indicate that HNRNPU is required to sustain a proliferative, RNA-processing-competent state in human cells: its loss slows expansion and shifts the transcriptome toward a p53/apoptotic stress program, and the spliceosome and nuclear-chromosome gene sets are among those depleted.

### *HNRNPU* loss rewires alternative splicing, predominantly exon skipping

Next, we analyzed alternative splicing with rMATS across the five standard event classes. Of 245,885 events tested, 36,277 reached an FDR below 0.05 and 21,728 also passed the effect-size threshold of an absolute dPSI of at least 0.1 (**Figure 3a and Table S2**). Skipped-exon events were by far the most abundant class (153,388 tested; 13,627 high-confidence events), followed by high-confidence events in mutually exclusive exons (3,555), retained introns (1,782), alternative 3’ splice sites (1,435) and alternative 5’ splice sites (1,329).

**Figure 3.**
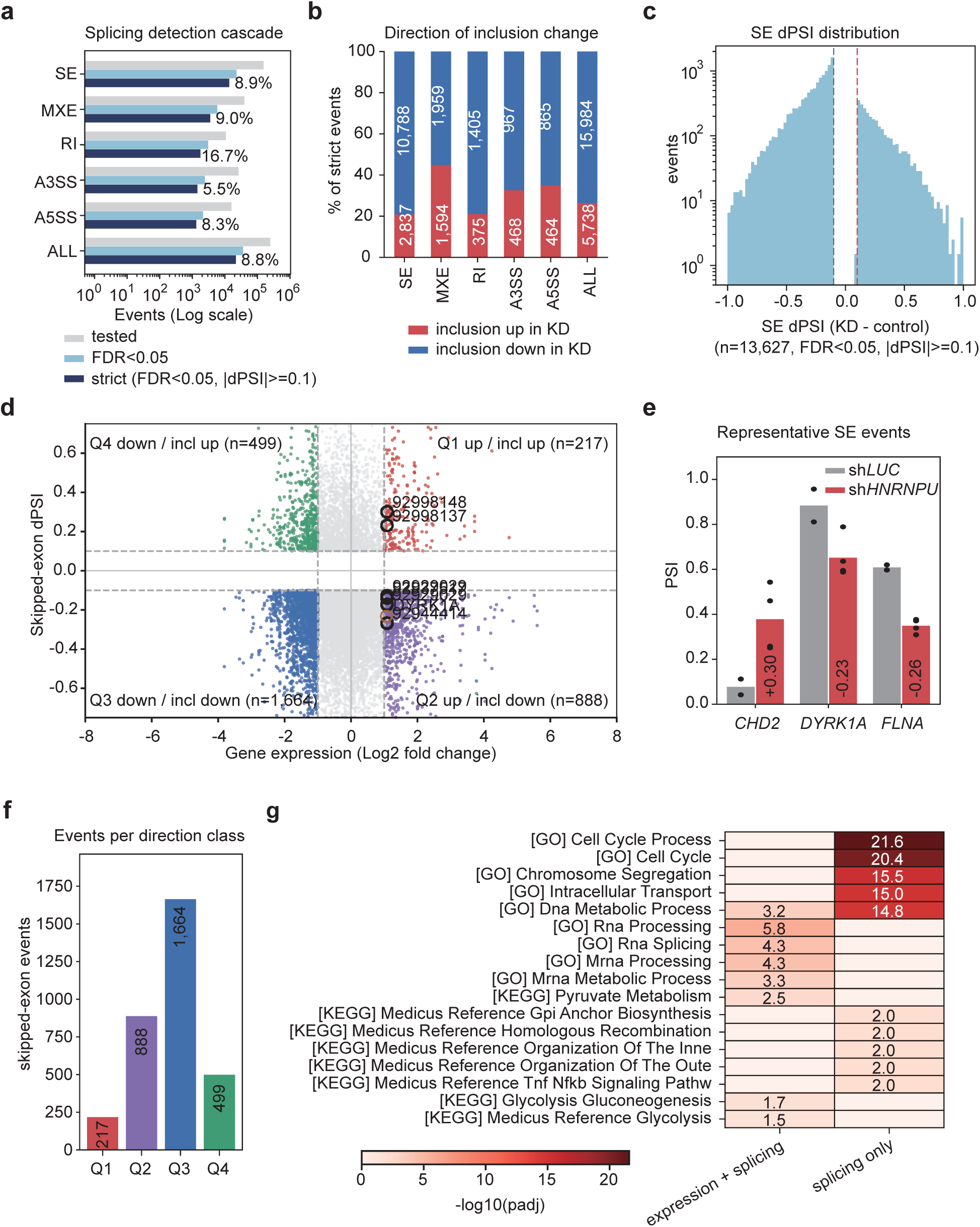
*HNRNPU* loss rewires alternative splicing, predominantly exon skipping. a. Numbers of tested, FDR-significant and high-confidence (FDR < 0.05, absolute dPSI of at least 0.1) events for each of the five rMATS event classes. b. Composition of inclusion-direction change among high-confidence events. c. Distribution of dPSI for high-confidence skipped-exon events. d. Gene expression change plotted against the dPSI of each individual high-confidence skipped-exon event, one point per event (n = 13,074); the four direction classes are defined per event, and the six *CHD2* events and the single *DYRK1A* event are circled. e. Number of skipped-exon events in each direction class. f. PSI of representative skipped-exon events in control and knockdown cells; individual replicate values are shown as dots. g. Pathway enrichment of genes altered at both the expression and the splicing level and of genes with a splicing change only, tested against all detected genes (gene ontology biological process and KEGG, Benjamini-Hochberg adjusted P < 0.05); the color scale shows minus log10 adjusted P and the value is printed in each cell.

The direction of change was markedly asymmetric. Among high-confidence skipped-exon events, 2,837 showed increased inclusion whereas 10,788 showed decreased inclusion (**Figure 3b**), and the global dPSI distribution was shifted toward negative values (**Figure 3c**). A bias of this kind is typical of cells in which a core splicing factor is limiting ^[19]^ and is consistent with a role for *HNRNPU* in exon recognition ^[7]^, although we did not measure the strength of the affected splice sites.

We next asked whether these splicing changes were accompanied by changes in transcript abundance. Because a single gene can harbor multiple events that move in opposite directions (*CHD2* itself spans −0.27 to +0.30, **Table S3**), we scored direction event by event rather than from a gene-level mean. Of the 13,074 high-confidence skipped-exon events with matched expression data, 3,268 lay in genes that also passed the differential-expression threshold and were distributed across all four quadrants: 1,664 showed reduced expression with reduced inclusion, 888 increased expression with reduced inclusion, 499 reduced expression with increased inclusion, and 217 increased expression with increased inclusion (**Figure 3d and 3e**). The six *CHD2* events and the single *DYRK1A* event are marked in **Figure 3d**. The remaining 9,806 events lay in genes whose expression change did not reach the threshold, indicating that most splicing changes were not accompanied by a substantial change in transcript abundance. Among the events highlighted in **Figure 3f**, *HNRNPU* depletion increased exon inclusion (PSI) in *CHD2* and decreased it in *DYRK1A* and *FLNA*.

Enrichment analysis of the two classes, defined by whether expression also changed, gave distinct functional signatures (**Figure 3g**). Genes altered at both levels were enriched for RNA processing (adjusted *P* = 1.7 × 10^-6^), RNA splicing (4.5 × 10^-5^) and mRNA processing (5.3 × 10^-5^), whereas genes with a splicing change but no expression change were enriched for cell-cycle processes (adjusted *P* = 2.6 × 10^-22^), chromosome segregation (3.2 × 10^-16^), intracellular transport (9.8 × 10^-16^) and DNA repair (1.5 × 10^-15^). Splicing changes in genes that also change in abundance therefore fall preferentially on the RNA-processing machinery itself, whereas abundance-independent splicing changes preferentially affect proliferation and genome-maintenance genes.

The more informative feature of these data is not that splicing changed, but how the changes were distributed. Roughly more than half of the affected events (9,806 of 13,074) lay in genes whose mRNA abundance did not pass the differential-expression threshold, so for most of the transcriptome the consequence of HNRNPU loss is a change in which transcript is produced rather than in how much of it is produced. This decouples the isoform layer from the abundance layer and means that the functional output of *HNRNPU* loss cannot be inferred from expression profiling alone: a gene may appear unchanged while its coding potential, domain composition or stability is altered. The minority of events that did coincide with expression changes were not a random sample; they fell preferentially on RNA-processing genes, whereas abundance-independent events affected cell-cycle and genome-maintenance genes. This partitioning suggests that *HNRNPU* loss erodes the same biological programs along two axes at once — depleting transcripts of the processing machinery while changing the structure of transcripts that maintain proliferation and genome integrity — and that the two effects can reinforce each other without either appearing as a single expression signature. Abundance-based readouts of *HNRNPU* loss therefore risk understating its functional consequences.

### Orthogonal evidence converges on *CHD2* as a neurodevelopmental node

To relate splicing changes to changes in transcript abundance, we intersected the genes carrying high-confidence skipped-exon events with the differentially expressed genes. Of 5,522 genes carrying at least one high-confidence skipped-exon event and 5,643 differentially expressed genes, 1,222 genes were altered at both levels (**Figure 4a**).

**Figure 4.**
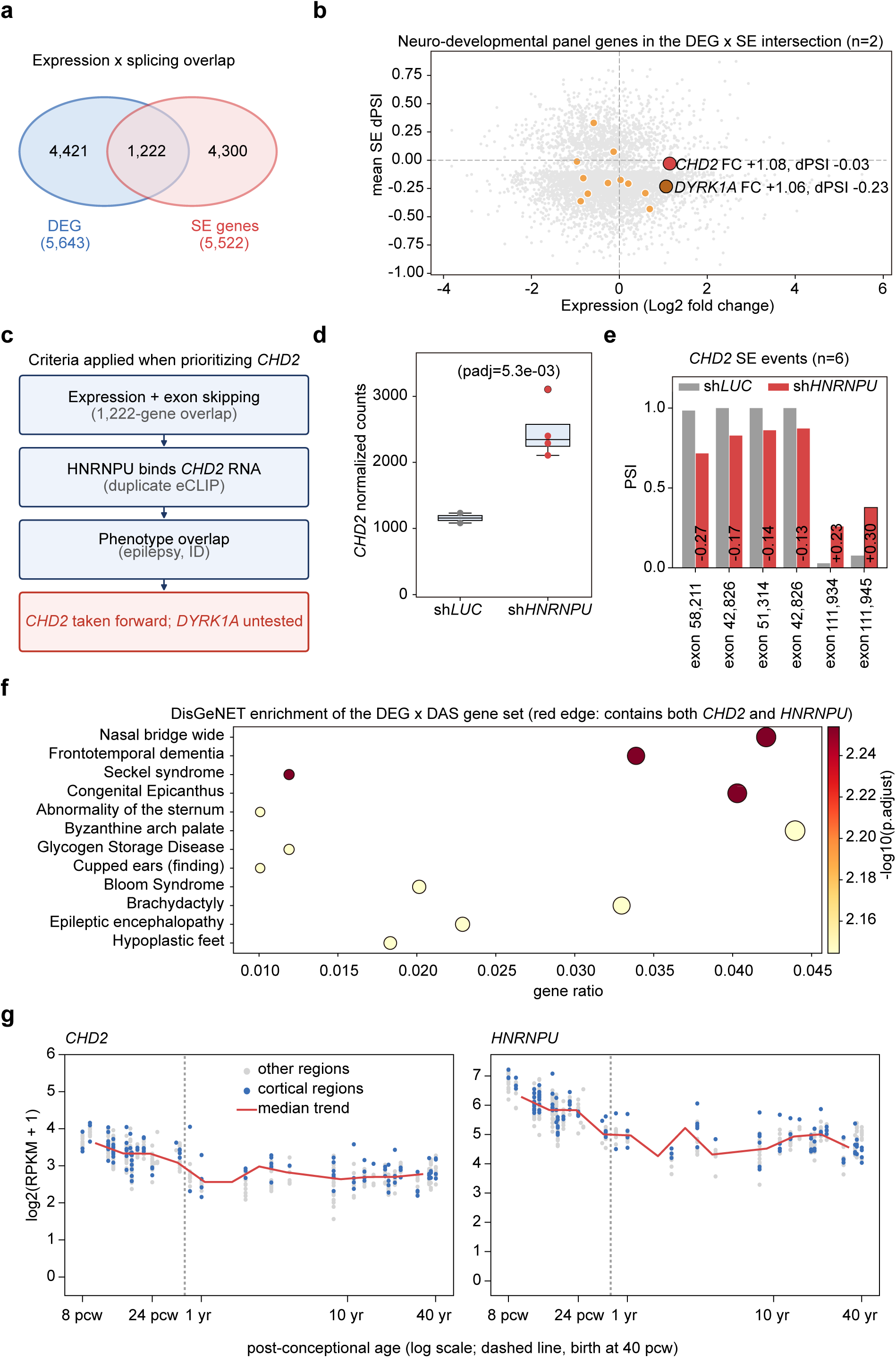
Orthogonal evidence converges on *CHD2* as a neurodevelopmental node. a. Overlap between *HNRNPU*-knockdown differentially expressed genes and genes carrying a high-confidence skipped-exon event. b. Neurodevelopmental panel genes plotted by expression change and mean dPSI; *CHD2* and *DYRK1A* are highlighted. c. Schematic of the criteria applied when prioritizing *CHD2*, and of the two candidates that the expression x splicing intersection contained. d. *CHD2* mRNA abundance in control and *HNRNPU* knockdown cells (normalized counts). e. PSI of the six *CHD2* skipped-exon events in control and knockdown cells; the dPSI and FDR of each event are given in Table S3. f. DisGeNET enrichment of the expression × splicing gene set; point size indicates gene count, color indicates significance, and outlined points indicate terms containing both *CHD2* and *HNRNPU*. g. *CHD2* and *HNRNPU* expression across human brain development (BrainSpan RNA sequencing atlas, 8 post-conceptional weeks to 40 years, 524 samples); each point is one sample, blue points are cortical regions and the red line is the median of each age bin.

We next asked which of the 1,222 genes altered at both levels were already implicated in a neurodevelopmental phenotype, because a pre-existing disease annotation makes a gene a more plausible mediator of the *HNRNPU* phenotype than one without. We therefore annotated the intersection against a panel of 32 genes that covers the phenotypic axes of *HNRNPU* syndrome — developmental and epileptic encephalopathy, intellectual disability, autism spectrum disorder and cortical malformation — as well as the molecular classes through which *HNRNPU* may act: chromatin and transcriptional regulation, synaptic adhesion and scaffolding, and the neuronal cytoskeleton (**Table S4**). The panel served as a prior-annotation lookup rather than as a statistical filter. Thirteen of its 32 genes carried at least one high-confidence skipped-exon event, and two of these, *CHD2* and *DYRK1A*, were also differentially expressed; we therefore took both forward as candidate nodes (**Figure 4a-c**).

*CHD2* was among the most strongly affected of these candidates. *CHD2* mRNA was increased in *HNRNPU*-depleted cells (log2 fold change +1.08, adjusted *P* = 5.3 × 10^-3^; **Figure 4d**), and six high-confidence exon-skipping events mapping to four distinct *CHD2* exons were detected, with dPSI values between −0.27 and +0.30 (**Figure 4e**). Four of the six events, involving canonical exons 5, 6 and 10, had lower inclusion in the knockdown (dPSI −0.27 to −0.13), whereas the two events involving the alternative exon 30a had higher inclusion (dPSI +0.23 and +0.30), so *HNRNPU* loss changes which *CHD2* exons are included rather than simply reducing *CHD2* output.

Because an intersection-based approach is susceptible to selection bias, we then asked which of the 1,222 genes already carry a neurodevelopmental annotation in the literature. Disease enrichment of the overlap between the expression & splicing gene set returned 43 terms at an adjusted P < 0.05, dominated by neurodevelopmental and epilepsy-related phenotypes, and *CHD2* and *HNRNPU* co-occurred as member genes in the most significant terms (**Figure 4f**). Two points limit how much this step can carry. The enrichment was computed on the same gene set that produced the candidates, so it is supporting context rather than independent evidence, and the terms are broad, the co-occurrence of two genes within them is weak evidence on its own.

The same panel identified a second candidate, *DYRK1A*, which also changed at both levels: its expression change was almost identical to that of *CHD2* (log2 fold change +1.06, adjusted P = 2.3 × 10^-3^), and it carried one high-confidence skipped-exon event (86 bp, dPSI −0.23, FDR 0.023). *DYRK1A* is itself required for neural progenitor proliferation, neuronal migration and synapse function ^[40]^, and it is dosage-sensitive in the human brain ^[41]^, it is best regarded as a second candidate node rather than as a contrast to *CHD2*. Two quantitative comparisons, both of which we state with their limits, separate the candidates in our data. First, the burden of regulated events: 6 of the 29 skipped-exon events tested in *CHD2* reached significance (21%), against a genome-wide rate of 8.9% (odds ratio 2.7, P = 0.039), whereas only 1 of 9 tested *DYRK1A* skipped-exon events did (11%, P = 0.57), so *CHD2* carries a modest excess of regulated exons while *DYRK1A* does not differ detectably from the background; the two genes are not, however, distinguishable from each other at these counts (Fisher exact P = 1.0). Second, co-expression with *HNRNPU* across brain development: *DYRK1A* ranks well below *CHD2* (**Table S8**). Peak occupancy does not separate the genes, because the peak density means that the absence of a call from the 86 bp *DYRK1A* exon is expected at these densities. The skipped *DYRK1A* exon is also 86 bp and therefore frame-shifting, so *DYRK1A* remains a plausible second node; we took *CHD2* forward because it combines the clinical overlap, the excess of regulated exons and the co-expression behaviour, and we do not claim that our transcriptomic data select it uniquely.

To ask whether these genes are expressed at the developmental stages at which the phenotype appears, we used the BrainSpan RNA sequencing atlas, which covers 524 samples from 8 post-conceptional weeks to 40 years ^[38]^. Both genes were expressed throughout development, with only a modest decline after birth: the median log2(RPKM + 1) fell from 3.43 in fetal samples to 2.70 in adults for *CHD2*, and from 5.94 to 4.79 for *HNRNPU* (**Figure 4g**). *HNRNPU* was the more highly expressed of the two at every stage. We next asked how closely each of the two candidates follows *HNRNPU* across these samples, because co-expression across brain development is one of the few independent criteria available for choosing between them: *CHD2* tracked *HNRNPU* closely (Pearson r = 0.84, 97th percentile of the 16,256 expressed genes), whereas *DYRK1A* did not (r = 0.67, 84th percentile; **Table S8**). This comparison is descriptive, because co-expression in bulk tissue reflects shared expression programs rather than direct regulation, and it is one of two criteria that separate the candidates in our data, the other being the excess of regulated exons.

### *HNRNPU* occupies *CHD2* pre-mRNA and constrains a frameshift-prone exon-skipping switch

To test whether the relationship could be direct, we asked whether *HNRNPU* occupies *CHD2* transcripts in vivo, using a publicly available duplicate *HNRNPU* eCLIP experiment performed in K562 cells (ENCSR520BZQ; duplicate 1, ENCFF408HQB; duplicate 2, ENCFF638WUB). Within the *CHD2*-201 transcription unit the two duplicates contained 184 and 220 peak calls; 107 calls in duplicate 1 overlapped a call in duplicate 2 (104 in the reciprocal comparison), and these 107 shared peaks were taken as the reproducible set (**Figure 5a**). The zoomed views in **Figure 5b** show that both duplicates contain peaks in the introns flanking each of the four regulated cassette exons (**Table S5**), and the inclusion changes of these exons are quantified in **Figure 5c**. Calling was dense across the whole transcription unit rather than focused on the regulated exons, consistent with binding along the length of the transcript rather than at a discrete site. Intra-exonic calls were sparse and inconsistent between duplicates (one call in a single replicate for canonical exons 6 and 10, one in each replicate for the alternative exon 30a, and none for exon 5), which is expected at the observed peak density and was not used to rank the exons.

**Figure 5.**
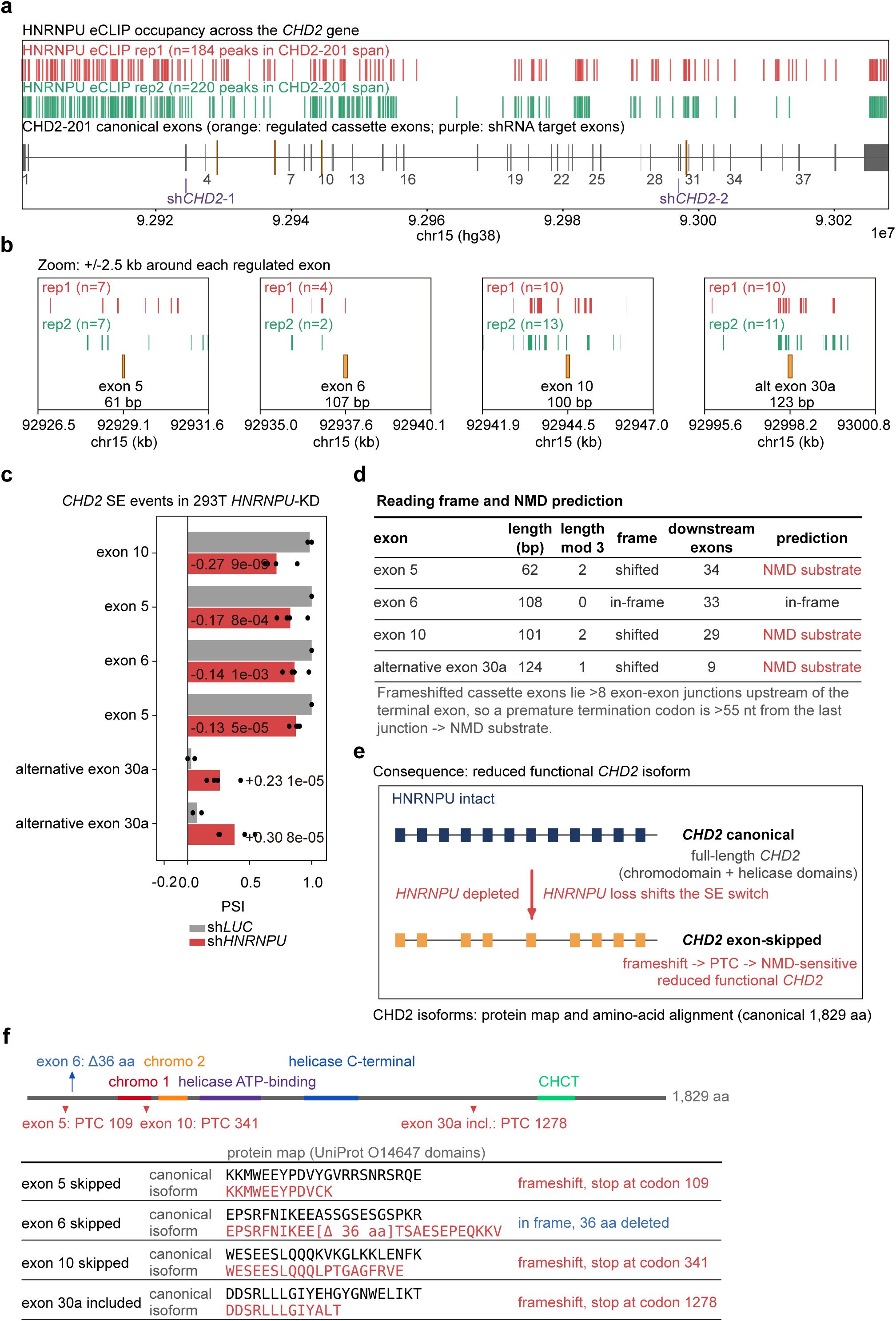
*HNRNPU* occupies *CHD2* pre-mRNA and constrains a frameshift-prone exon-skipping switch. a. *HNRNPU* eCLIP peak calls from the two duplicates of a single K562 experiment (ENCSR520BZQ; 184 and 220 calls, of which 107 were present in both) across the *CHD2*-201 transcription unit, with the 39 canonical exons shown below; orange exons are the regulated cassette exons and purple boxes mark the exons targeted by the shRNAs used in this study. b. Occupancy in windows of plus or minus 2.5 kb around each regulated exon, shown for the two duplicates (dup1, dup2); the number above each track is the number of eCLIP peaks in that window and the exon length is given beneath each panel; because calls are dense across the whole transcription unit (one per 0.6-0.7 kb), these counts are descriptive and do not by themselves show preferential binding at the regulated exons. c. PSI of the six *CHD2* skipped-exon events after *HNRNPU* depletion, with dPSI and FDR given beneath each event. d. Reading-frame and nonsense-mediated decay prediction for the four unique regulated exons: exon length, length modulo three, frame consequence, number of downstream canonical exons and predicted outcome; full coordinates and distances to the stop codon are given in Table S3. e. Schematic of the proposed consequence, in which *HNRNPU* loss shifts *CHD2* exon inclusion toward frameshift-bearing transcripts that are predicted to be degraded by nonsense-mediated decay; the outcome shown assumes that the frameshift transcripts are degraded by NMD. a. Protein consequence of the regulated cassette exons: domain map of canonical *CHD2* (UniProt O14647; chromodomains 1 and 2, helicase ATP-binding and helicase C-terminal domains, and the CHCT domain) with the premature stop codon of each frameshift isoform marked below the chain, followed by the amino-acid alignment of the canonical protein and each isoform around the first divergent codon.

To ask what these inclusion changes would do to the protein, we first asked whether the four exons preserve the reading frame (**Figure 5d**). Exon 6 is 108 bp, a multiple of three, so its skipping removes 36 residues without shifting the reading frame; the other three exons (62, 101 and 124 bp) are not multiples of three, so their skipping or inclusion shifts the frame. In each of these three cases the resulting premature stop codon lies 1.4–4.7 kb upstream of the last exon-exon junction of the spliced transcript, far beyond the 50–55 nucleotide rule, and 30–99 kb from the annotated termination codon in genomic terms (**Table S3**), so the transcripts are predicted substrates of nonsense-mediated decay ^[37]^. If these transcripts are degraded, the expected consequence of *HNRNPU* depletion is a reduced supply of full-length *CHD2* rather than a new protein species (**Figure 5e**). Consistent with this prediction, the alternative exon 30a is annotated only in a *CHD2* transcript that Ensembl itself classifies as a nonsense-mediated decay biotype (*CHD2*-216).

We then translated each affected isoform to locate the truncation relative to the protein domains (**Figure 5f**). Starting from the MANE Select transcript (*CHD2*-201, ENST00000394196; 5,487 nt of coding sequence encoding 1,829 amino acids, with no internal stop codon), we removed each canonical exon or inserted the alternative exon, translated the resulting open reading frame and compared the predicted proteins with the canonical sequence. Skipping of exon 5 truncates the protein after 108 residues (stop at codon 109), skipping of exon 10 after 340 residues (codon 341), and inclusion of the alternative exon 30a after 1,277 residues (codon 1,278). The exon-5 and exon-10 products lack both the chromodomain and the helicase domain, whereas the exon-6 product is an in-frame deletion of 36 residues.

To ask whether these exons are already covered by the known variant spectrum, we intersected them, their core splice sites and 2, 10 and 50 bp of flanking sequence with the 1,968 *CHD2* short variants in ClinVar that carry a GRCh38 position ^[39]^. No ClinVar variant falls within any of the four exons, their splice sites or their immediate flanks, and the closest reported variant lies 1.1 kb from the alternative exon 30a. The regulated exons are therefore not established variant hot spots, which is consistent with a mechanism that acts on exon inclusion rather than through coding variation. We therefore propose that *HNRNPU* depletion shifts *CHD2* exon inclusion toward frameshift-bearing, NMD-sensitive isoforms. Two outcomes are then possible, and our data do not distinguish them: if the frameshift transcripts are degraded efficiently, the functional isoform must account for the rise in total *CHD2* mRNA, whereas if they escape or saturate NMD they accumulate and the rise in total mRNA is their signature. Isoform-resolved measurement is therefore required to decide between them.

### *CHD2* expression tracks *HNRNPU* dosage across public transcriptomes

To ask whether the relationship could be observed independently of our knockdown, we first examined *HNRNPU* occupancy across the *CHD2* transcription unit in the public data (**Figure 6a**). Both replicates contained peaks within 2.5 kb of all four regulated exons, but occupancy was not exon-selective. The two duplicates contained 184 and 220 peak calls across the approximately 130-kb *CHD2*-201 interval, equivalent to one call per 0.6-0.7 kb, and 22 of the 39 canonical exons contained at least one peak within 250 bp. The four regulated exons and their 250-bp flanks comprise 1.9% of the transcription unit but contained 16 of the 404 calls, only 2.1-fold their proportional share. All four regulated windows were occupied in both replicates, but the density and broad distribution of the calls do not establish preferential binding. These data therefore support an association between *HNRNPU* and *CHD2* transcripts without localizing a functionally relevant binding site.

**Figure 6.**
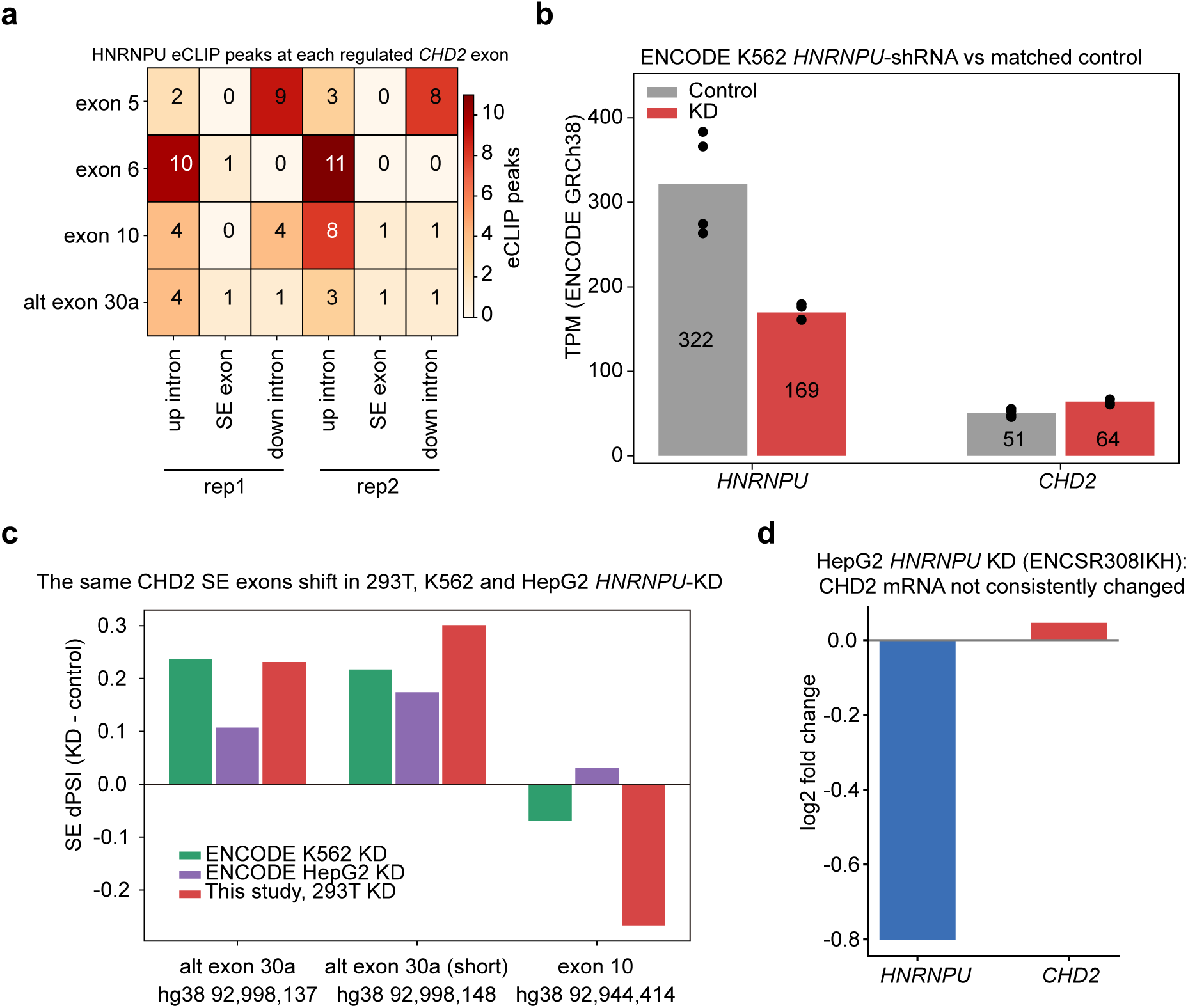
Public datasets reproduce the *HNRNPU-CHD2* relationship. b. Number of *HNRNPU* eCLIP peak calls at the introns flanking, and within, each regulated *CHD2* exon, for both duplicates. c. *HNRNPU* and *CHD2* TPM in the K562 *HNRNPU* shRNA experiment and its matched non-targeting control; points are individual quantification sets and bars are means. d. dPSI of corresponding *CHD2* skipped-exon events in HEK293T cells (this study), K562 and HepG2. e. *HNRNPU* and *CHD2* expression change in the HepG2 *HNRNPU* knockdown (ENCSR308IKH).

The ENCODE K562 *HNRNPU* shRNA experiment provided an independent replication. Compared with its matched non-targeting control, *HNRNPU* mRNA fell from 321.8 to 169.5 TPM (log2 ratio −0.93), showing that the knockdown was effective, while *CHD2* mRNA rose from 50.6 to 64.1 TPM (log2 ratio +0.34; **Figure 6b**); the expression of the *CHD2* change is smaller than the +1.08 seen in HEK293T cells, as expected for a partial knockdown in a different cellular context. The alternative exon 30a changed in the same direction as in our data (hg19 chr15:93,541,367-93,541,491, corresponding to hg38 chr15:92,998,137-92,998,261; dPSI +0.237, FDR 9.0 × 10^-5^; **Figure 6c**). Across the 5,021 genes carrying high-confidence skipped-exon events in both datasets, the mean dPSI per gene correlated between the two studies (Spearman ρ = 0.11, P = 2.4 × 10^-14^) and the direction of change agreed for 59% of genes (binomial P = 1 × 10^-40^); among the genes that reached significance in K562 the agreement was higher (n = 762, ρ = 0.29, 67% concordant). The public knockdown therefore reproduces the direction of the *HNRNPU*-dependent splicing program, although only weakly at the genome-wide level, and with smaller effect sizes that are consistent with the shallower depletion in that experiment.

The HepG2 data provided only partial and internally inconsistent evidence for replication. In the differential-expression file associated with a detectable *HNRNPU* knockdown, *HNRNPU* decreased by log2 fold change −0.80, whereas *CHD2* was essentially unchanged (+0.05, adjusted P = 0.98); the other available quantifications varied in both the magnitude and direction of the *HNRNPU* effect and were not treated as independent replication. At the splicing level, the HepG2 estimate for exon 10 was opposite in sign to our result but negligible and non-significant (+0.031, FDR = 1.0; normalized estimate +0.009), whereas the short form of alternative exon 30a was concordant in one quantification (+0.174, FDR = 0.0038) but not after normalization (+0.097, FDR = 1.0) (**Figure 6c and 6d**). These results are compatible with cell-type-dependent regulation, but the available data cannot distinguish biological context from differences in knockdown efficiency or quantification.

### *CHD2* loss perturbs cell-cycle and neural gene programs in SH-SY5Y cells

Finally, to ask whether *CHD2* has functional consequences relevant to a neural context, we depleted *CHD2* in the human neuroblastoma line SH-SY5Y using shRNAs. The two shRNAs reduced *CHD2* mRNA efficiently (shCHD2-1, log2 fold change −0.86, adjusted *P* = 4.6 × 10^-13^; shCHD2-2, −0.90, adjusted P = 7.5 × 10^-14^, **Figure 7a**), and joint analysis of the two effective shRNAs identified 479 differentially expressed genes (**Figure 7b**), of which 288 changed in the same direction under both shRNAs (**Table S6**). Pathway analysis of the downregulated genes showed enrichment for cell cycle, homologous recombination and Fanconi anemia (**Figure 7c**), consistent with the established role of *CHD2* in chromatin remodeling and genome maintenance. Aggregated phenotype scoring confirmed coordinated downregulation of proliferation and DNA-repair gene sets, with a smaller positive score for neural markers (**Figure 7d**).

**Figure 7.**
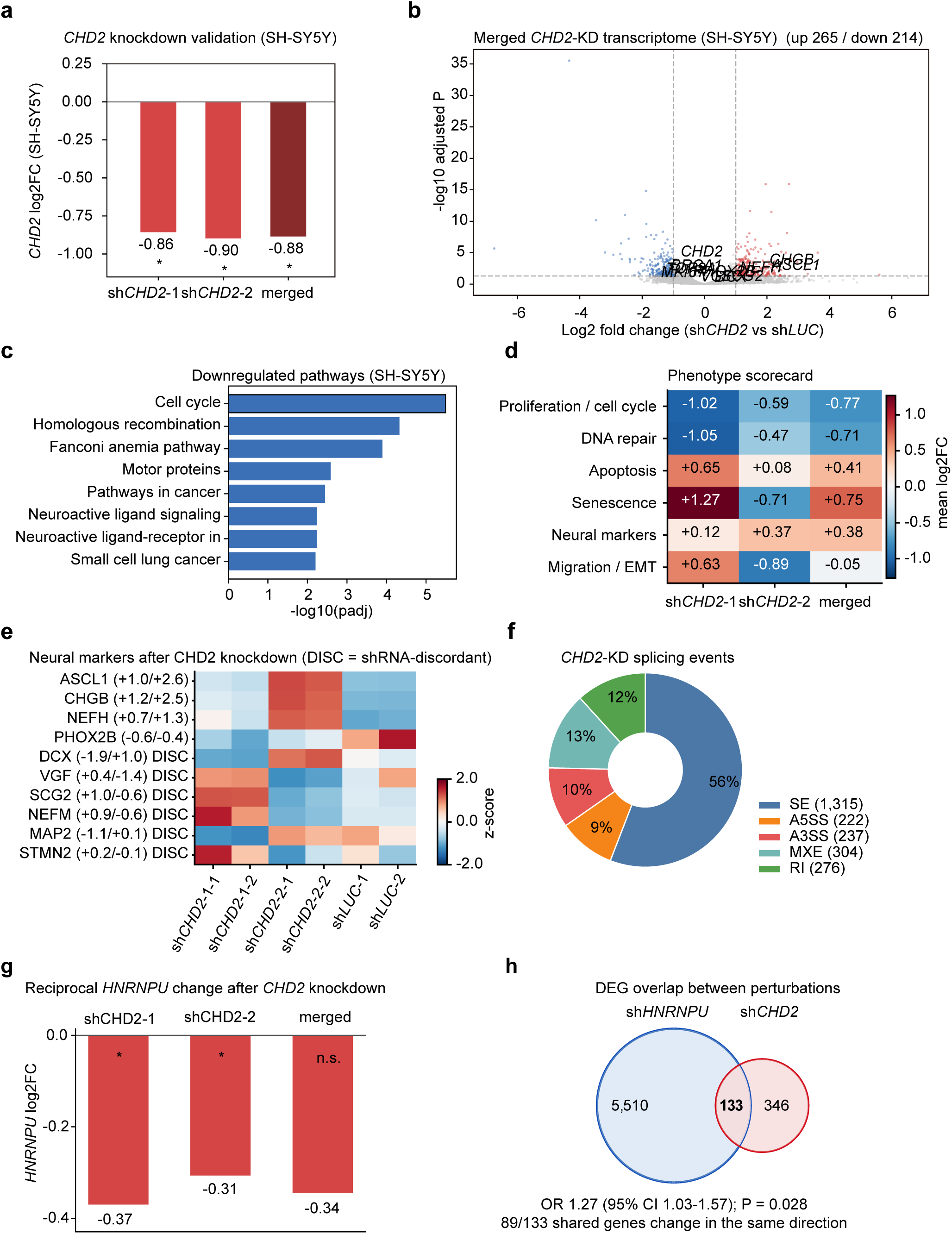
*CHD2* loss perturbs cell-cycle and neural gene programs in SH-SY5Y cells. a. *CHD2* knockdown efficiency for the two effective shRNAs and for the merged shCHD2-1 plus shCHD2-2 comparison; shCHD2-A did not reduce *CHD2* and was excluded from the panels below. b. Volcano plot of the merged *CHD2* knockdown. c. Downregulated KEGG pathways (adjusted P < 0.05). d. Phenotype scorecard showing the mean log2 fold change of curated gene panels under shCHD2-1, shCHD2-2 and the merged comparison. e. Expression of neural markers; values in parentheses are log2 fold changes under shCHD2-1 and shCHD2-2, and "DISC" marks markers whose direction differed between the two shRNAs. f. Composition of high-confidence splicing events after *CHD2* depletion (SE 1,315; A5SS 222; A3SS 237; MXE 304; RI 276). g. Reciprocal change in *HNRNPU* mRNA after *CHD2* depletion in SH-SY5Y cells, shown for the two effective shRNAs and the merged comparison; asterisks indicate adjusted P < 0.05 and n.s. indicates not significant. h. Overlap between HNRNPU-knockdown differentially expressed genes in HEK293T cells (5,643 genes) and *CHD2*-knockdown differentially expressed genes in SH-SY5Y cells (479 genes). Of the 16,629 genes quantified in both experiments, 133 were shared (odds ratio 1.27, 95% confidence interval 1.03-1.57; Fisher exact P = 0.028), and 89 of the 133 shared genes changed in the same direction.

Because shRNA experiments are vulnerable to seed-mediated off-target effects, we tested whether the gene sets used here were enriched for transcripts matching the 6-nucleotide or 7-nucleotide seed of either shRNA (**Table S6**). We applied this test to the *CHD2* knockdown, where the gene sets are small and curated and a seed effect could plausibly create an apparent signal; in the *HNRNPU* knockdown the equivalent control is the requirement that genes change in the same direction under two shRNAs with unrelated seed sequences. Across the twelve tests, none reached significance after correction for multiple testing (minimum adjusted *P* = 0.061); the strongest nominal signal was for the sh*CHD2*-1 7-nucleotide seed within the core consistent gene set (odds ratio 1.34, nominal *P* = 0.037, adjusted *P* = 0.148). The core consistent gene set is therefore unlikely to be explained by seed-mediated off-target repression, and the requirement that genes change in the same direction under two shRNAs with different seed sequences provides an additional, sequence-independent filter. The gene sets unique to each shRNA, which we excluded from the main analysis, did show nominal enrichment for seed-matched transcripts in two of four comparisons, supporting the decision to treat them separately.

Few neural markers changed consistently (**Figure 7e**). *ASCL1*, *CHGB* and *NEFH* were upregulated under both shRNAs, whereas *PHOX2B* was downregulated; several other markers, including *DCX*, *SCG2*, *VGF*, *NEFM* and *MAP2*, changed in opposite directions under the two shRNAs and were therefore not used as evidence. *CHD2* depletion also altered splicing, with 1,315 significant skipped-exon events in the merged analysis (**Figure 7f**).

We also examined the reciprocal direction. *HNRNPU* mRNA was modestly lower after *CHD2* depletion (sh*CHD2*-1, log2 fold change −0.37, adjusted *P* = 4.2 × 10−3; sh*CHD2*-2, −0.31, adjusted *P* = 0.035; merged, −0.34, adjusted *P* = 0.31), a weak and inconsistent signal compatible with feedback but insufficient to establish a reciprocal arm (**Figure 7g**). The two perturbations shared 133 differentially expressed genes (odds ratio 1.27, *P* = 0.028), representing 2.4% of the genes regulated by *HNRNPU* (**Figure 7h**). An overlap of this size would be expected if *CHD2* accounts for a small part of the *HNRNPU*-dependent program, but it is equally consistent with the shared growth-arrest response of the two knockdowns. We therefore describe *CHD2* as a candidate downstream node rather than as a phenocopy of *HNRNPU* loss (**Figure 7g and 7h**).

## Discussion

We report that *HNRNPU* depletion changes both the abundance and the splicing of *CHD2*, that *HNRNPU* occupies *CHD2* pre-mRNA in an independent cell line at positions flanking exactly the exons that change, and that translating the affected isoforms predicts truncated CHD2 proteins rather than a simple loss of transcript: skipping of exon 5, skipping of exon 10 and inclusion of the alternative exon 30a each introduce a premature termination codon at codons 109, 341 and 1,278, whereas skipping of exon 6 removes 36 residues in frame. Together with the observation that *CHD2* loss alters cell-cycle, genome-maintenance and neural gene programs, these data outline a candidate *HNRNPU-CHD2* axis that connects *HNRNPU* dosage to a neurodevelopmental gene program.

*HNRNPU* has been reported to influence splicing globally through U2 snRNP maturation ^[7]^, and our data are consistent with this: exon skipping was the dominant event class and the direction of change was strongly biased toward reduced inclusion. This asymmetry is more consistent with a role for *HNRNPU* in the inclusion of a subset of exons than with a uniform loss of splicing activity. An alternative interpretation is that *HNRNPU* loss reduces the fidelity of exon definition, with poorly defined exons affected preferentially. Our data do not distinguish between these possibilities, because we did not measure splice-site strength; a minigene or in vitro splicing system would be required to do so.

A difficulty for a simple model in which *HNRNPU* activates *CHD2* is that *CHD2* mRNA increases rather than decreases after *HNRNPU* depletion. Our data suggest a resolution: if *HNRNPU* constrains the inclusion of exons whose skipping is frameshift-inducing, then loss of *HNRNPU* would be expected to produce more, but less functional, *CHD2* transcript, with the increase in total mRNA reflecting either compensatory transcription or reduced NMD efficiency. This isoform-balance interpretation makes a specific and testable prediction, namely that the ratio of functional to frameshift *CHD2* isoform should fall after *HNRNPU* depletion even as total *CHD2* mRNA rises. Testing it requires isoform-resolved quantification, for example long-read sequencing, exon-junction-specific quantitative PCR or N- and C-terminal antibodies, none of which were available here.

What matters for *CHD2* function is also where these changes fall in the protein: skipping of exon 5 truncates the protein before the chromodomain and skipping of exon 10 truncates it within the chromodomain, so neither product retains the helicase domain, whereas the alternative exon 30a is inserted downstream of the helicase domains. If these transcripts are degraded by nonsense-mediated decay, the net effect of *HNRNPU* loss would be a reduced supply of full-length *CHD2* rather than a new protein species, which is the behavior expected of a chromatin remodeler that must be present at sufficient dosage during cortical neurogenesis ^[13, 14]^. Splicing changes of this kind are not only pathogenic but also therapeutically tractable, as shown by the modulation of *SMN2* exon 7 inclusion in spinal muscular atrophy ^[42]^. Two observations support the relevance of this axis to neurodevelopment. First, both genes are dosage-sensitive in the human brain: heterozygous loss of either causes epilepsy with developmental delay ^[8, 9, 15–17]^, and mouse models of reduced *HNRNPU* dosage show cortical and behavioral deficits ^[10, 11]^. Second, *CHD2* depletion in a neural cell line changed cell-cycle and DNA-repair programs and altered the expression of a small number of neural markers, which is compatible with a chromatin remodeler acting in neural cells, although cell-cycle and DNA-repair signatures of this kind also accompany impaired proliferation. The prediction that follows is testable in patient material: if the axis operates during human brain development, *HNRNPU* haploinsufficiency should reduce full-length *CHD2* transcript and protein in neural cells, and the two disorders should share a component of their transcriptional phenotype.

*CHD2* is a SNF2-family chromatin remodeler that is highly expressed in the developing brain and is required for normal neurodevelopment ^[15–17]^. The pathways downregulated after *CHD2* depletion in SH-SY5Y cells (cell cycle, homologous recombination, Fanconi anemia) are consistent with a role in chromatin and genome maintenance, and the accompanying changes in neural markers are consistent with a requirement for *CHD2* in neural cells, but they rest on a small number of markers, several of which changed in opposite directions under the two shRNAs and were therefore not used as evidence. These observations support the plausibility of *CHD2* as a functional node downstream of *HNRNPU*, but they do not demonstrate mediation.

Because the discovery space was defined by the intersection of two analyses, the risk that the prioritized gene is an arbitrary member of a large set deserves explicit attention, and we therefore summarize the prioritization criteria used here in **Figure 4c**. We should be explicit about what this prioritization can and cannot show. The panel is not a statistical filter: two of its 32 members entered the intersection, which is close to the number expected from the size of the intersection itself, and both of the two candidates are established neurodevelopmental disease genes with phenotypes that overlap that of *HNRNPU* syndrome ^[8, 9, 15–17, 41]^, so the panel identifies candidates rather than selecting among them. The occupancy data show binding to the *CHD2* transcript but, at the densities observed, do not localize the binding or demonstrate that the regulated exons are preferred. The functional evidence is coherent with *CHD2* biology but was obtained in a different cell line and cannot exclude a contribution from the growth arrest that accompanies both knockdowns. What the data do support is that *CHD2* is one of at least two plausible nodes in the intersection, that its exon inclusion changes in a way predicted to truncate the protein, and that its loss perturbs cell-cycle and neural programs; *DYRK1A*, which was identified by the same intersection, remains a second candidate that was not functionally tested here, rather than a rejected one.

Two independent lines of public evidence converge on the same relationship. *HNRNPU* eCLIP places the protein on *CHD2* pre-mRNA at the regulated exons, with peaks present in both duplicates at all four, and this measurement does not depend on the efficiency of any knockdown. The K562 knockdown experiment, in a different cell type and with a shallower depletion, reproduces the direction of the *CHD2* expression change, the change in the same skipped exon and, genome-wide, the direction of the *HNRNPU*-dependent splicing program (**Figure 6b and 6c**), with an *HNRNPU* reduction that is demonstrable in the same samples. We therefore regard the public data as reproducing the relationship in one of the two cell types tested; the causal step, and the question of which exons are functional targets, remain to be tested experimentally.

### Several limitations qualify these conclusions

The discovery dataset (*HNRNPU* depletion in HEK293T cells) and the neural evaluation dataset (*CHD2* depletion in SH-SY5Y cells) come from different cell lines, so the design cannot establish a same-cell causal chain. The reciprocal experiment provided limited support for feedback: *HNRNPU* fell modestly after each effective *CHD2* shRNA but not in the merged comparison (**Figure 7g**), and 133 differentially expressed genes, representing 2.4% of the *HNRNPU*-regulated genes, were shared between the two perturbations (**Figure 7h**). This overlap is compatible with *CHD2* mediating a small component of the *HNRNPU*-dependent program, but it is also consistent with a shared growth-arrest response and does not establish that *CHD2* depletion phenocopies *HNRNPU* loss. Protein-level confirmation is lacking: the reading-frame and nonsense-mediated decay predictions follow from translation of the affected isoforms and the established 50-55 nucleotide rule ^[37]^, but they were not validated with isoform-specific assays, and no N- or C-terminal antibodies were available. The occupancy evidence is also indirect. It was measured in K562 rather than in HEK293T or SH-SY5Y cells, it rests on duplicate-level peak calls because the ENCODE IDR-thresholded set for that experiment (180 peaks genome-wide) contains no *CHD2* peak, and binding does not by itself prove regulation. The public perturbation data are likewise limited in magnitude and cell-type dependence: the K562 knockdown was incomplete and the *CHD2* change there is about one third of that in HEK293T cells, and the genome-wide concordance with our data is weak (Spearman ρ = 0.11, 59% of genes in the same direction), whereas HepG2 provided only partial and internally inconsistent evidence and did not show a *CHD2* expression change in the analysis with a clear *HNRNPU* knockdown. Three further caveats follow from the numbers rather than from the design. The occupancy calls are dense, so co-localization with the regulated exons does not by itself show preferential binding, and the panel of neurodevelopmental genes is a source of candidates rather than a statistical filter, since two of its members entered an intersection, whose size predicts a similar number; both knockdowns impair proliferation, so cell-cycle and genome-maintenance signatures cannot be attributed to a specific function of the depleted gene. Finally, the *HNRNPU* knockdown used two biological replicates per group and a merged two-shRNA design, and the disease-enrichment analysis was performed on a gene set derived from the same data that motivated the hypothesis, so the disease annotation is supporting context rather than independent evidence.

The strategy used here is general and does not depend on the specific genes involved. Intersecting differential expression with differential splicing, then requiring an independent disease annotation and an independent occupancy dataset, produces a short candidate list that is robust to the main sources of bias in each individual analysis. The same framework could be applied to other RNA-binding proteins implicated in neurodevelopmental disorders ^[19]^, particularly where a knockdown produces modest expression changes but widespread splicing changes. Two design choices were important in our hands: analyzing two shRNAs jointly so that candidate genes must survive a sequence-independent filter, and using publicly available occupancy and perturbation data to distinguish binding from consequence. Both choices are available to any laboratory that has performed a knockdown followed by RNA sequencing, and both are compatible with the growing collection of ENCODE and related datasets.

### Conclusions and future directions

*HNRNPU* occupies *CHD2* pre-mRNA, and its depletion is accompanied by a reorganization of *CHD2* exon inclusion, with three of four regulated exons predicted to generate NMD-sensitive, frameshift-bearing transcripts; *CHD2* loss in a neural cell line perturbs cell-cycle and neural programs. The most informative next experiments are, in our view: isoform-resolved measurement of *CHD2* transcripts after *HNRNPU* depletion, using long-read sequencing or exon-junction-specific assays together with N- and C-terminal antibodies; rescue of *HNRNPU*-depletion phenotypes by expression of an shRNA-resistant *CHD2* cDNA, ideally in the same cell line; a temporally resolved, same-cell-line bidirectional experiment to determine whether the relationship is unidirectional or contains a feedback component; and validation in patient-derived neurons carrying *HNRNPU* loss-of-function variants.

## Author contributions

H.T.: investigation, formal analysis, visualization, writing - original draft. Z.X.: investigation, formal analysis. R.W.: investigation. S.H.: resources, writing - review and editing. I.A.B.: formal analysis, writing - review and editing. A.P.H.: supervision, writing - review and editing. G.M.: conceptualization, supervision, funding acquisition, writing - original draft, writing - review and editing. All authors reviewed and approved the final version of the manuscript.

## Declaration of generative AI and AI-assisted technologies in the writing process

During the preparation of this work the authors used ChatGPT (GPT-5, OpenAI) to assist with language editing. After using this tool, the authors reviewed and edited the content and take full responsibility for the content of the publication. All figures were produced from the primary data with standard analysis and plotting software and were assembled in Adobe Illustrator; no generative AI tool was used to create, alter or enhance any figure or image.

## Ethics approval and consent to participate

This study used only the established human cell lines HEK293T and SH-SY5Y. No human participants, human tissue or animals were involved, and no personal data were analyzed, so ethical approval and informed consent were not required.

## Data availability

The HEK293T *HNRNPU* knockdown and SH-SY5Y *CHD2* knockdown RNA sequencing data generated in this study will be deposited in the Gene Expression Omnibus (GEO). The public datasets analyzed here are available from the ENCODE portal: *HNRNPU* eCLIP in K562 (ENCSR520BZQ), *HNRNPU* shRNA in K562 (ENCSR047IUS) with its matched non-targeting control (ENCSR438MDN), and *HNRNPU* shRNA in HepG2 (ENCSR308IKH). Human brain expression data were obtained from the BrainSpan Atlas of the Developing Human Brain (brainspan.org). Analysis code and derived tables are provided with the manuscript and are available from the corresponding author on request.

## Supplementary Materials

**Table S1.** Differential expression after *HNRNPU* depletion in HEK293T cells (merged knockdown versus shLUC; limma).

**Table S2.** High-confidence alternative-splicing events after *HNRNPU* depletion, with event counts by class.

**Table S3.** Regulated *CHD2* skipped-exon events with coordinates, PSI, dPSI and FDR.

**Table S4.** Neurodevelopmental panel of 32 genes with expression and splicing statistics that led to the intersection.

**Table S5.** *HNRNPU* eCLIP peaks mapped to *CHD2* exons and flanking introns.

**Table S6.** *CHD2* knockdown in SH-SY5Y: differential expression, core consistent genes, phenotype panel scores and off-target seed tests.

**Table S7.** Datasets analyzed in this study: cell line, assay, replicates, source accession and use for the two experiments generated here and the four public ENCODE datasets.

**Table S8.** BrainSpan co-expression of *CHD2*, *DYRK1A* and reference genes with *HNRNPU* across 524 samples.

